# Mapping of Neuropeptide Y Expression in *Octopus* Brains

**DOI:** 10.1101/2020.04.24.056465

**Authors:** Gabrielle C. Winters, Gianluca Polese, Anna Di Cosmo, Leonid L. Moroz

**Affiliations:** Whitney Laboratory for Marine Biosciences, University of Florida, St. Augustine, FL, 32080, USA; Di Cosmo Laboratory, University of Napoli Federico II, Department of Biology, Napoli, Italy; Departments of Neuroscience and McKnight Brain Institute, University of Florida, Gainesville, FL, 32610, USA

**Author notes:** Correspondence: Leonid L. Moroz The Whitney Laboratory, 9505 Ocean Shore Blvd, St Augustine FL, 32086, USA Anna Di Cosmo.

**Keywords:** neuropeptide, cephalopod, *Nautilus*, nervous system evolution, feeding, reproduction

## Abstract

Neuropeptide Y (NPY) is an evolutionarily conserved neurosecretory molecule implicated in a diverse complement of functions across taxa and in regulating feeding behavior and reproductive maturation in *Octopus*. However, little is known about the precise molecular circuitry of NPY-mediated behaviors and physiological processes, which likely involve a complex interaction of multiple signal molecules in specific brain regions. Here we examined the expression of NPY throughout the *Octopus* central nervous system. The sequence analysis of *Octopus* NPY precursor confirmed the presence of both signal peptide and putative active peptides, which are highly conserved across bilaterians. *In situ* hybridization revealed distinct expression of NPY in specialized compartments, including potential “integration centers,” where visual, tactile, and other behavioral circuitries converge. These centers integrating separate circuits may maintain and modulate learning and memory or other behaviors not yet attributed to NPY-dependent modulation in *Octopus*. Extrasomatic localization of NPY mRNA in the neurites of specific neuron populations in the brain suggests a potential demand for immediate translation at synapses and a crucial temporal role for NPY in these cell populations. We also verified the presence of NPY mRNA in a small cell population in the olfactory lobe, which is a component of the *Octopus* feeding and reproductive control centers. However, the molecular mapping of NPY expression only partially overlapped with that produced by immunohistochemistry in previous studies. Our study provides a precise molecular map of NPY mRNA expression that can be used to design and test future hypotheses about molecular signaling in various *Octopus* behaviors.

**Research Highlights/Graphical Abstract text:** Neuropeptide Y (NPY), an evolutionarily conserved neurosecretory molecule, is expressed in specialized regions of the *Octopus* brain controlling feeding, reproduction, and visual and tactile memory circuits. Extrasomatic mRNAs were found in neurites, suggesting synaptic synthesis of NPY.

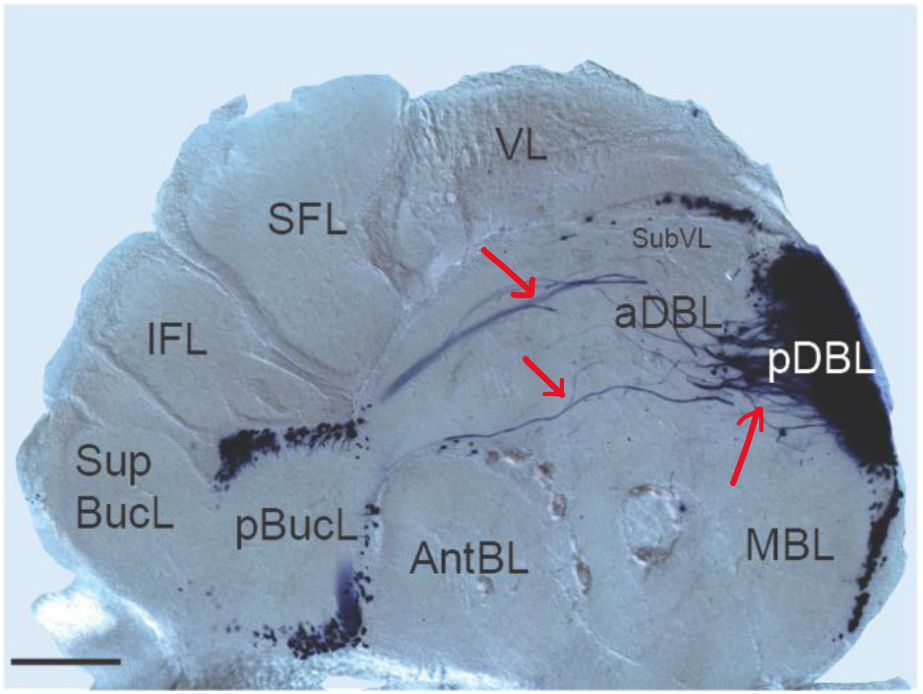

## 1 Introduction

A diverse array of functions have been attributed to the 36-amino acid polypeptide Neuropeptide Y (NPY) (Thorsell *et al.*, 2001) since its initial discovery in mammals in 1982 (Tatemoto K Fau - Carlquist *et al.*, 1982), including digestive (Cox, 1993), cognitive (Redrobe *et al.*), and cardiovascular (Walker *et al.*, 1991) processes. Several interdisciplinary studies have revealed shared functions of NPY in controlling metabolic states (Ruffin *et al.*, 1997, Beck, 2006, Holzer *et al.*, 2012, Salomäki-Myftari *et al.*, 2016, Fadda *et al.*, 2019), circadian rhythms (Albers *et al.*, 1984, Yannielli *et al.*, 2001, Erion *et al.*, 2016) and stress/anxiety (Wahlestedt *et al.*, 1993, Thorsell *et al.*, 2002, Wu *et al.*, 2003) responses across diverse taxa, but whether these functional commonalities are the result of evolutionary conservation or independent recruitment is yet unknown.

The role of NPY in the regulation of feeding and satiety has been explored in both protostome and deuterostome taxa since the 1990s (Hanson *et al.*, 1995). One extensively conserved role for NPY is as an orexigenic (appetite-stimulating) factor (Beck, 2006, Christie *et al.*, 2011, Di Cristo *et al.*, 2017, Fadda *et al.*, 2019). This has been examined closely in the fly *Drosophila melanogaster*, where Neuropeptide F (*D. melanogaster* NPY ortholog) neurons were shown to regulate many aspects of feeding, including olfactory learning (Rohwedder *et al.*, 2015) and temporal foraging behavioral shifts (Wu *et al.*, 2003) in larvae, motivational control of appetitive memory in the mushroom bodies of adults (Krashes *et al.*, 2009), and central modulation of peripheral olfactory responses (Lee *et al.*, 2017). NPF/NPY expression has also been associated with diminished stress, fear, and anxiety responses in both mammals (Wahlestedt *et al.*, 1993, Thorsell *et al.*, 2002) and fruit flies (Wu *et al.*, 2003). NPF in fruit flies also mediates clock-regulated sex dimorphic behavior (Lee *et al.*, 2006) and circadian gene expression (Erion *et al.*, 2016).

Neuropeptide Y orthologs have been identified and functionally characterized in representatives from three classes of the phylum Mollusca: Cephalopoda, Bivalvia, and Gastropoda. Injection of purified NPY (aka NPF) ligand into the saltwater Manila clam (a bivalve) *Ruditapes philippinarum*, increased filtration rates, which presumably increased nutrient acquisition (Wang *et al.*, 2017). The roles role for NPY homologs in gastropods appears to vary by taxa. In the pond snail, *Lymnaea stagnalis*, administration of NPY appeared to inhibit both growth and reproduction but had no discernible effect on the intake of food (de Jong-Brink *et al.*, 2001). Neuropeptide Y injections in the gastropod *Aplysia californica* surprisingly reduced food intake, and its activity appears to be a key element in the feeding network reconfiguration as the animal’s motivational state shifts from hunger to satiety (Jing *et al.*, 2007). Therefore, the role of NPY in molluscs is not entirely conserved but consistently plays some role in mediating the acquisition and/or digestion of nutrients.

One proposed function of NPY in *Octopus* is as a candidate messenger mediating the temporal shift between growth and reproductive life stages, which is an essential part of their ecological life history (Di Cosmo *et al.*, 2014, Polese *et al.*, 2015). The transition from hunting and feeding behaviors to *Octopus* sexual maturity is irreversible in nature (Wodinsky, 1977, Boyle, 1983); it is hormone-controlled, and must only occur once the individual’s lifetime nutritional needs have been met (De Lisa *et al.*, 2012, Di Cosmo *et al.*, 2014, Polese *et al.*, 2015). Gametogenesis and subsequent reproductive processes are energetically demanding in females, and in many cases controlled by an array of hormones and other secretory molecules that initiate senescence and, ultimately, death (Boyle, 1983, De Lisa *et al.*, 2012, Di Cosmo *et al.*, 2013, Di Cosmo *et al.*, 2016). Upon sexual maturation, the metabolic requirements of the reproductive system overtake the demands of somatic tissues (Di Cosmo *et al.*, 2013). Only after an animal accumulates sufficient nutrient reserves to meet reproductive demands do specialized signals to growth and reproduction neural centers induce a shift from the feeding state to the reproductive life stage (Wodinsky, 1977, Boyle, 1983).

The primary CNS structures composing the growth and reproduction control centers are the olfactory lobes^1^ (Di Cosmo *et al.*, 1998, Di Cosmo *et al.*, 2014, Polese *et al.*, 2015) and optic glands (Wells *et al.*, 1959, Wells, 1978, Juárez *et al.*, 2019, Wang *et al.*, 2019), both located between the brain and the hilum of each optic lobe on the optic stalks. The subpedunculate lobes, located in the posterior/dorsal region below the subvertical lobe, are also involved in the central control of these physiological processes. A current model (Di Cristo, 2013, Di Cosmo *et al.*, 2014, Polese *et al.*, 2015) for feeding and reproduction regulation in *Octopus* implies recruitment of multiple neuropeptides and neurohormones including FMRFamide (Di Cosmo *et al.*, 1998, Suzuki *et al.*, 2002), Gonadotropin-releasing hormone (GnRH) (Di Cosmo *et al.*, 1998, Di Cristo *et al.*, 2009), Galanin (Suzuki *et al.*, 2000) and Neuropeptide Y (Suzuki *et al.*, 2002) (NPY). Briefly, this model suggests that the olfactory lobes receive information about an individual’s energy stores/demands, and this input determines whether neurons in the olfactory lobe produce Neuropeptide Y (NPY): a primary candidate messenger for optic gland inhibition. To date, the mechanism by which the NPY-producing neurons of the olfactory lobe receive satiation input is still unknown. According to this model, if the individual has not accumulated needed energy reserves, NPY is constitutively produced by the olfactory lobe, maintaining the animal’s appetite so *Octopus* can continue to feed and grow. Once lifetime energy demands are met, Neuropeptide Y production ceases in the olfactory lobe, and the olfactory lobe peptides GnRH and Galanin activate the optic gland (Di Cosmo *et al.*, 2014); the animal can enter the reproductive life stage. The optic gland then produces a yet undescribed trophic factor(s) that can activate the process of gonadal maturation and the onset of mating behaviors. Thus, the specific roles and, in particular, localized expression of neuropeptides and neurohormones are critical to understanding the mechanistic control of feeding and reproduction.

Here, we have focused on exploring the region-specific expression and distribution of Neuropeptide Y in *Octopus*. By cloning and characterizing the *Octopus* NPY precursor, we mapped the expression of mRNA transcripts encoding neuropeptide Y prohormone (pNPY-the precursor to the active peptide NPY) throughout the entire central nervous system. These experiments have provided molecular insights to existing models for mediation of feeding and reproduction, as well as presented novel information that can be used to generate hypotheses and future studies geared toward deciphering functional roles for NPY in *Octopus*.

## 2 Materials and Methods

We obtained the DNA sequence for *Octopus vulgaris Pro-neuropeptide Y* (*OvpNPY*) from our Illumina sequencing data (BioSample Accession number SAMN09698694) using tblastn (Basic Local Alignment Search Tool (Altschul *et al.*, 1990) using a protein query to search a DNA database) with the protein sequence from *Lymnaea stagnalis* (CAB63265.1) as a probe. The coding region of the sequence (flanked by sequences specific to T7 (Roche T7 RNA polymerase Sigma cat no 1088176700) and T3 (Roche T3 RNA polymerase-Sigma cat no 11031163001) promoters, and Not1 (NEB cat no R3189S) and Pme1 (NEB cat no R0560S) restriction enzymes) was synthesized into a pUC57 plasmid DNA vector by Genscript®.

### Animal Acquisition and Preparation

Due to the limited availability of *Octopus vulgaris*, we elected to use the readily available species *Octopus bimaculoides* for expression localization studies. Nucleotide sequences for ObpNPY and OvpNPY share 97% identity, so probes generated using the synthesized OvpNPY sequence were sufficiently similar and specific in both species.

Six wild-caught adults >50 g *Octopus bimaculoides* were shipped overnight to our laboratory from Marinus Scientific in Long Beach, CA, in Summer 2016. Octopuses were anesthetized with either 337 mM MgCl_2_, 10% ethanol (EtOH) in the filtered seawater (FSW) on ice, or placed directly on ice for euthanasia. Neuronal tissues (Brain, Optic Lobes, Arm Cords, and Stellate Ganglia were extracted and fixed in 4% paraformaldehyde (PFA) in FSW for subsequent *in situ* hybridization.

The experiments in the present study were conducted in accordance with the principles and procedures that were approved by the Institutional Animal Care of University on Naples Federico II (Project n° 608/2016-PR-17/06/2016; protocol n. DGSAF 0022292-P-03/10/2017), and according to the Italian and European law (European Directive 2010/63 EU L276; Italian DL. 4/03/2014, no. 26; the ethical principles of Reduction, Refinement, and Replacement).

### Probe Preparation

All Digoxygenin (DIG) labeled probes were generated using the Roche DIG labeling kit (Sigma Cat no 11277073910). Antisense probes were generated using Pme1 (NEB cat no R0560S) restriction enzyme followed by T7 RNA polymerase (Sigma cat no 10881767001). Final probes were cleaned up using RNeasy MinElute Cleanup Kit (Qiagen Cat No./ID: 74204), and one microliter was visualized on a 2% agarose gel to estimate concentration.

### NPY *in situ* hybridization

The procedure for *in situ* hybridization was based on a modified protocol for *Aplysia californica* described previously (Jezzini *et al.*, 2005, Antonov *et al.*, 2007, Moroz *et al.*, 2013). Dissected neuronal tissues from six individual octopuses were removed whole and incubated in 4% paraformaldehyde (PFA) in phosphate-buffered saline (PBS) at 4°C for three hours. Tissues were then rinsed in PBS before slicing at 175 (brain) or 250 (other tissues) micrometer thickness on a vibratome. Slices were then fixed overnight in 4% PFA in phosphate-buffered saline (PBS) at 4°C. The following day slices were dehydrated stepwise and stored in 100% methanol until use (up to three weeks).

Next, dehydrated tissue slices were rehydrated stepwise and taken through a series of washes to optimize permeability. After prehybridization, probes were added at a concentration of 1 µg/μL in hybridization buffer, and tissues were incubated overnight at 50°C. Next, after a series of washes and blocking steps, tissues were incubated at 4°C in a solution of 0.05% alkaline phosphatase-conjugated DIG antibodies (Roche cat #11093274910) overnight. After a final series of washes, the tissues were incubated in 20 µL NBT/BCIP per 1 mL detection buffer. Upon development completion, the tissues were incubated in 4% PFA in MeOH for 40-60 minutes and washed twice in 100% EtOH before being stored in 100% EtOH or mounted on a slide using methyl salicylate and Permount.

### Microscopy and Imaging

Images of tissue preparations were taken with a Qimaging Retiga EXi Fast1394 digital camera mounted on a Nikon TE-2000E microscope using DIC settings using with NIS Elements software V4.3. Whole images were enhanced for clarity in Adobe Photoshop. Figures were created using Adobe Illustrator and Microsoft PowerPoint.

### Protein Sequence and Structural Analyses

General domain architecture for pNPY protein sequences was determined using the Simple Modular Architecture Research Tool (SMART) (Letunic *et al.*, 2015, Letunic *et al.*, 2018). Domain features not predicted by SMART (e.g., glycine residues to indicate amidation sites and cleavage sites) were identified visually from the protein sequence alignment. This alignment was created using default parameters in Muscle (Edgar, 2004). Selected sequences were identified using BLASTp (Altschul *et al.*, 1990) searches on NCBI. The molecular phylogenetic analysis was completed by using the Maximum Likelihood method based on the Whelan And Goldman model (Whelan *et al.*, 2001). We depicted an image of the tree with the greatest log likelihood (−1477.4407). The bootstrap value (the percentage iterations in which the clustered taxa were associated with one another) is shown next to each node, except for those for which the value was below 50. Neighbor-Join and BioNJ algorithms were applied to a matrix of pairwise distances estimated using a JTT model to generate initial trees for the heuristic search. Then topology with superior log likelihood value was selected. Branch lengths correspond to the number of substitutions per site. These evolutionary analyses were conducted in MEGA7(Kumar *et al.*, 2016). Figures were created using Adobe Creative Suite®.

## 3 Results

### Neuropeptide Y as one of few evolutionarily conserved neuropeptides in Bilateria

Genes encoding the precursor of neuropeptide Y were only identified in protostomes and deuterostomes; they are absent in non-bilaterians sequenced so far (Ctenophora, Porifera, Cnidaria, Placozoa). Multiple sequence alignments (Figure 1a) revealed amino acid sequences of the NPY prohormone are highly conserved across bilaterians, particularly in the putative active sites (bioactive peptide after posttranslational modifications like cleavage (by prohormone convertases) and amidation (by peptidylglycine alpha-amidating monooxygenase (PAM) enzyme). The sequence for the predicted bioactive peptide for *O. bimaculoides* NPY shares 86% identity with both cephalopod species *Nautilus pompilius* and *Doryteuthis pealei*, as well as 72% with that of the gastropod *Aplysia californica*, and 27% with that of *Homo sapiens*. Beyond the putative NPY active site, sequence conservation is relatively low in signal peptide domains and C-termini. The *O. bimaculoides* predicted bioactive NPY peptide is MLAPPNRPSEFRSPEELRKYLKALNE YYAIVGRPRF-amide.

**Figure 1.**
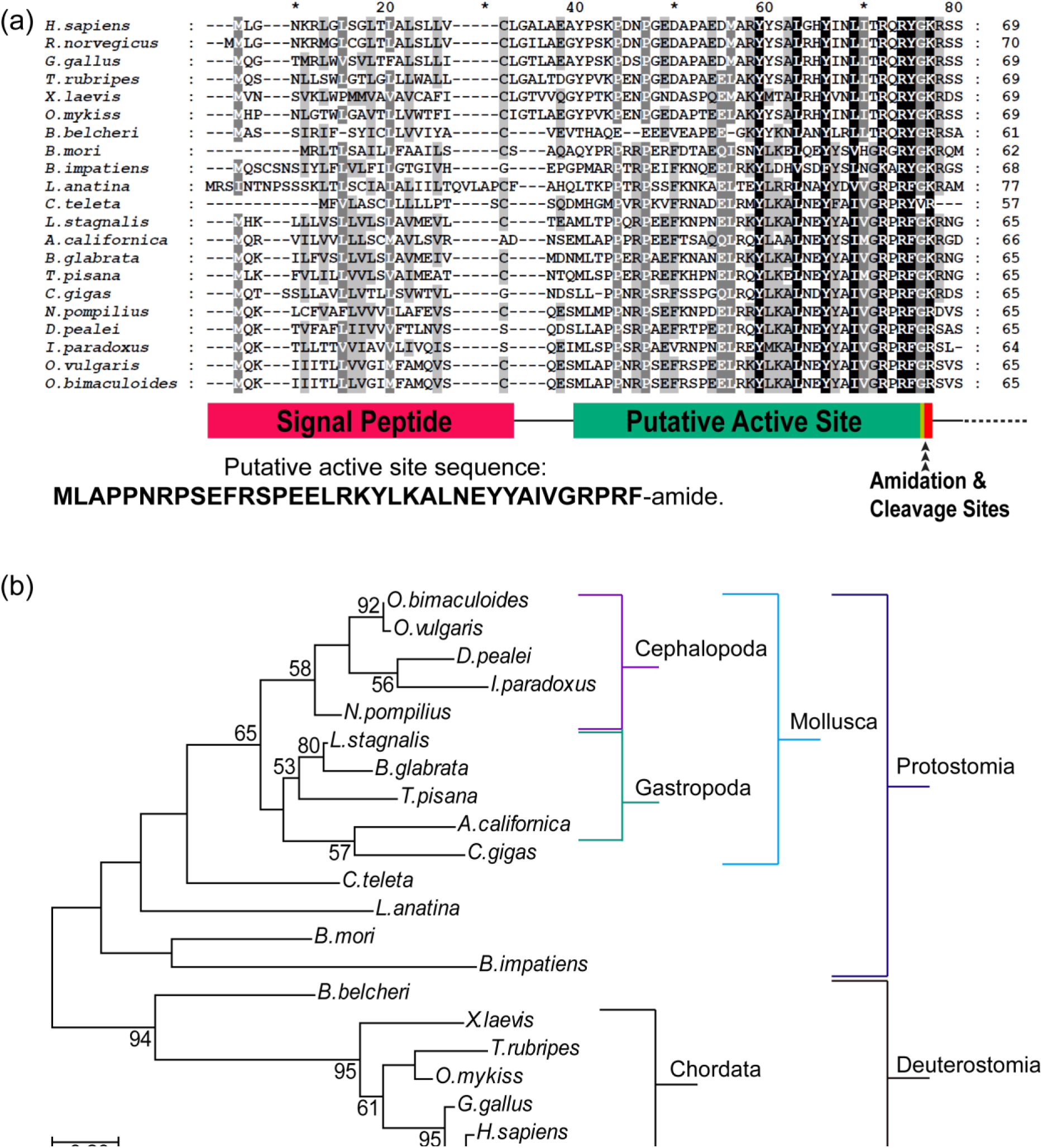
Multiple sequence alignment (a) and gene tree (b) for neuropeptide Y precursor (pNPY) protein sequences illustrate evolutionary conservation. (a) depicts the N-terminus of pNPY sequences with conserved residues aligned vertically using MUSCLE (Edgar, 2004) (note, entire sequences are not shown, as conservation is low or absent in the C-terminus beyond the predicted active site). Below, the alignment is a schematic of predicted domains of pNPY, based on predictions of SMART (Letunic *et al.*, 2015, Letunic *et al.*, 2018) software web tools and confirmed mammalian (Porcine) pNPY active site analyses (Tatemoto, 1982). Note that the glycine (G) at position 77 (thin yellow bar in the schematic) is followed by a dibasic cleavage site (KR) in the majority of taxa, but cephalopods exhibit only a monobasic cleavage site (R) in this position. Also note that in molluscs, the amino acid at position 76 is a phenylalanine (F) instead of the tyrosine (Y) for which the conserved polypeptide was originally named. (b) indicates a maximum likelihood gene tree generated using the Whelan And Goldman model (Whelan *et al.*, 2001). Taxon branches are denoted in brackets on the right. Branches with bootstrap support over 50% are noted, and branch lengths correspond to the number of substitutions per site. These evolutionary analyses were conducted in MEGA7 (Kumar *et al.*, 2016).

### Highly-localized expression of pNPY mRNA in the *Octopus bimaculoides* brain

NPY precursor-encoding transcripts (pNPY) appear to be expressed in very densely packed neurons of the dorsal basal lobe (DBL), subvertical lobes (SubVL), and in the moderately densely packed cell somata layer of the posterior buccal lobe. Thick projections extend up to 2 mm from the middle of the superior dorsal region of the DBL (Figure 2a) and are wide enough to be visible in a coronal cross-section through basal lobes below the vertical lobe (Figure 2d). Expression of pNPY encoding transcript appears absent in all other lobes except for a few large cells scattered at the periphery of the anterior basal lobe (Figure 2a).

**Figure 2.**
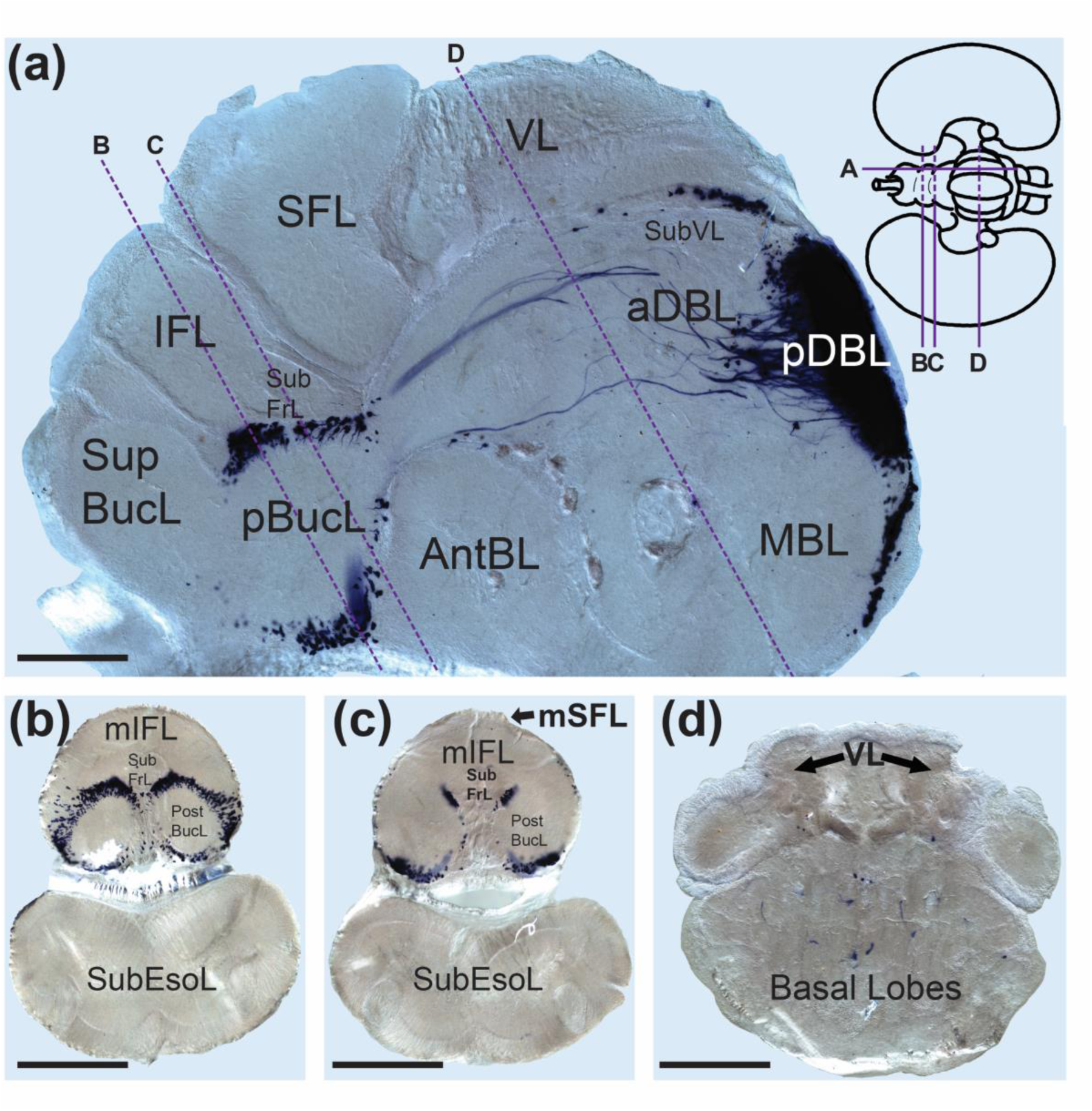
pNPY (neuropeptide Y encoding transcript) is expressed in multiple brain lobes in *O. bimaculoides*. (a) pNPY expression in a mediolateral brain slice in the sagittal plane. pNPY probe localized to the subvertical lobe (SubVL), dorsal basal lobe (DBL: aDBL is anterior, and pDBL is posterior), median basal lobe (MBL), posterior buccal lobe (pBucL), and in few cells in the superior periphery of the anterior basal lobe (AntBL). Some neurons of the dorsal basal lobe appear to have long thick processes that project through the basal lobe neuropil toward the frontal lobes. (b), (c), and (d) show brain slices in the diagonal coronal plane and are in order of most anterior to most posterior. Slices in (b) and (c) include both supra and sub-esophageal lobes and depict pNPY expression in the neuronal somata layer of the posterior buccal lobe (pBucL). (d) depicts only supraesophageal lobes and illustrates pNPY expression in cross-sections of the thick projections originating in the dorsal basal lobes. Scale bar measurements are (a) 500 μm, (b and c) 1 mm, and (d) 715 μm.

### pNPY is expressed in the Dorsal Basal and the Subvertical lobes

Seven morphologically distinct neuronal subtypes were identified (Table 1), expressing pNPY transcripts in the dorsal basal and subvertical lobes. Neurons with exceptionally long and thick pNPY-containing neurites originated in the middle of the superior dorsal region of the DBL (type 1) (Figures 2a, 3a and 3b), while neurons in the inferior/posterior cell somata layer of the DBL and median basal lobe (MBL) were relatively large (type 5; Figure 3f and 3g) and their processes are relatively short if present at all. The somata of all pNPY-containing neurons with the long thick processes (type 1) were clustered densely and appeared indistinguishable from one another, except for the neuronal soma indicated by the closed arrow in Figure 3b. Smaller neurons expressing pNPY are densely clustered in the subvertical lobe (type 6 in Figure 3e). Most of these neurons have no visible neurites recognized by pNPY probes.

**Table 1.**
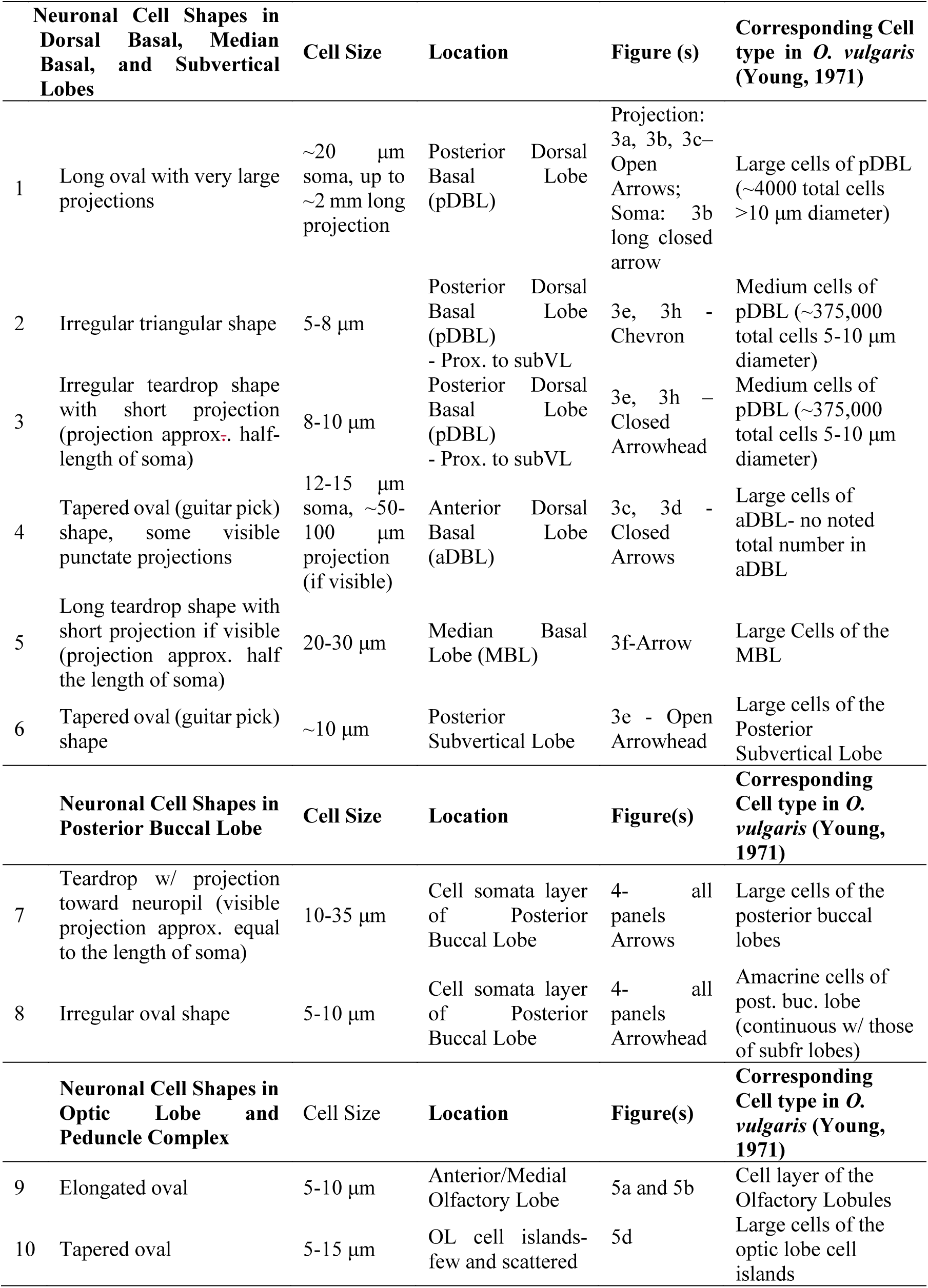
Cell morphology of pNPY transcript expressing neurons.

**Figure 3.**
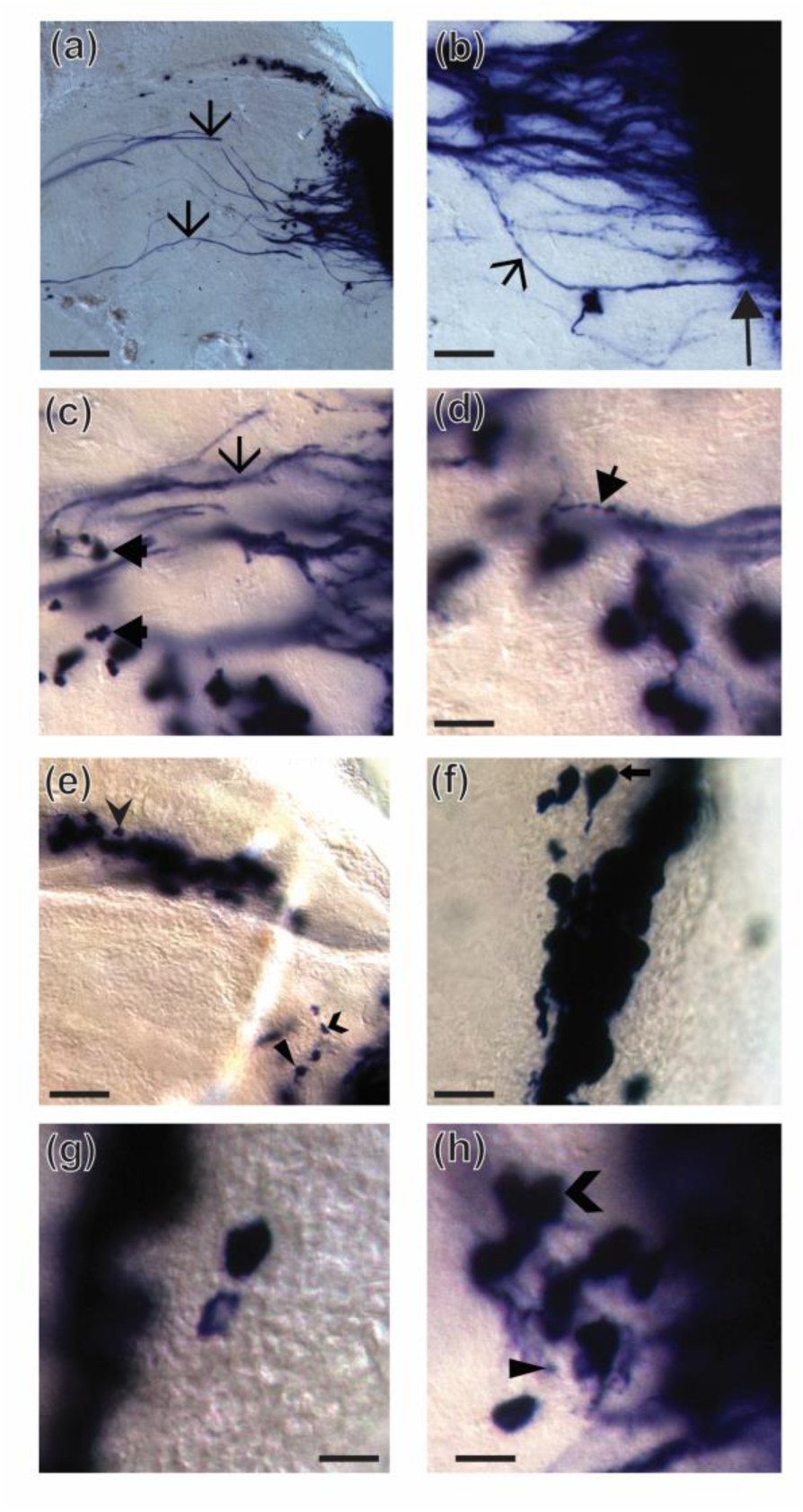
pNPY neurons of the posterior dorsal basal (DBL), median basal (MBL), and subvertical lobes (SubVL) exhibit morphological diversity. (a) through (d) highlight mRNAs of pNPY in axons projecting from the neurons of the DBL. The larger long projections (open arrows-type 1) appear to originate in the cell somata layer of the posterior DBL (pDBL) and can measure up to 2 mm in length and 20 µm in diameter. The long arrow in (b) denotes the oval soma of the neuron, with a neurite (short open arrow) projecting at least 600 µm into the pDBL neuropil toward the anterior DBL (aDBL). The closed arrows in panel (c) and (d) depict cell type 4- in (c), the closed arrows indicate the neuronal somata that appear to be in the wall of neurons delineating the anterior and posterior regions of the DBL. pNPY transcripts are also present in punctate patches of thin, shorter processes illustrated in panel (d) (closed short arrow-the projection of type 4). Panels (b) through (h) also highlight the morphological diversity of neuronal somata of cells expressing pNPY in the posterior region of the supraesophageal lobes. A total of six morphologically distinct neuronal subtypes were identified in the DBL, MBL, and SubVL. In (e) and (h), chevron arrowheads indicate type 2, and closed arrowheads indicate type 3. In (e), the open arrowhead indicates type 6 cells. The arrow in (f) indicates cell type 5. A summary of the neuronal somata diversity (pNPY neuron types one though six) in this region can be found in Table 1 with arrowhead identifier descriptions. Scale bar measurements are (a) 350 μm, (b) 100 μm, (c) 65 μm, (d &f) 35 μm, (e) 50 μm, (g &h) 20 μm.

The posterior region of the DBL (superior to the majority of the long projecting neurons (type 1), but inferior to the subvertical lobes) contains two distinct types of small neurons. The first, type 2, has an irregular triangle shape and no visible processes (indicated by the chevron-shaped arrow in Figures 3e and 3h), the second (type 3) has an irregular teardrop shape with short projections that are approximately half-length of the soma (indicated by the closed arrowhead arrow in Figures 3e and 3h). Amongst the large thick processes of pNPY neurons, type 4 are tapered oval (guitar pick) shaped neurons (indicated by thick line closed arrows in Figure 3c), some of which have visible punctate projections (indicated by thinner line closed arrows in Figure 3d).

### pNPY expression in the Posterior Buccal Lobe

Two distinct subtypes of neurons express pNPY encoding transcripts in the cell somata layer of the posterior buccal lobes (pBucL) (Table 1). These neurons are generally clustered densely together. Figure 4 shows the density and arrangement of pNPY positive cells. The pBucL neurons had a teardrop shape with kinked projections, typically pointing toward the neuropil that measures approximately equal to the length of their somata (Type 7 in table 1). These neurons exhibit a wide range of sizes (from 10 to 35 micrometers in soma length), as is depicted by the colored arrows in Figure 4b. In general, the larger neuronal somata were located toward the periphery of the cell somata layer, and the smaller ones were located toward the middle or (in fewer cases) adjacent to the neuropil. An additional class of smaller (type 8 in Table 1, 5-15 micrometer length cell somata) irregular oval-shaped neurons expressed pNPY (closed arrowheads in all panels of Figure 4). These cells appear to cluster toward the neuropil facing edge of the cell somata layer in the pBucL.

**Figure 4.**
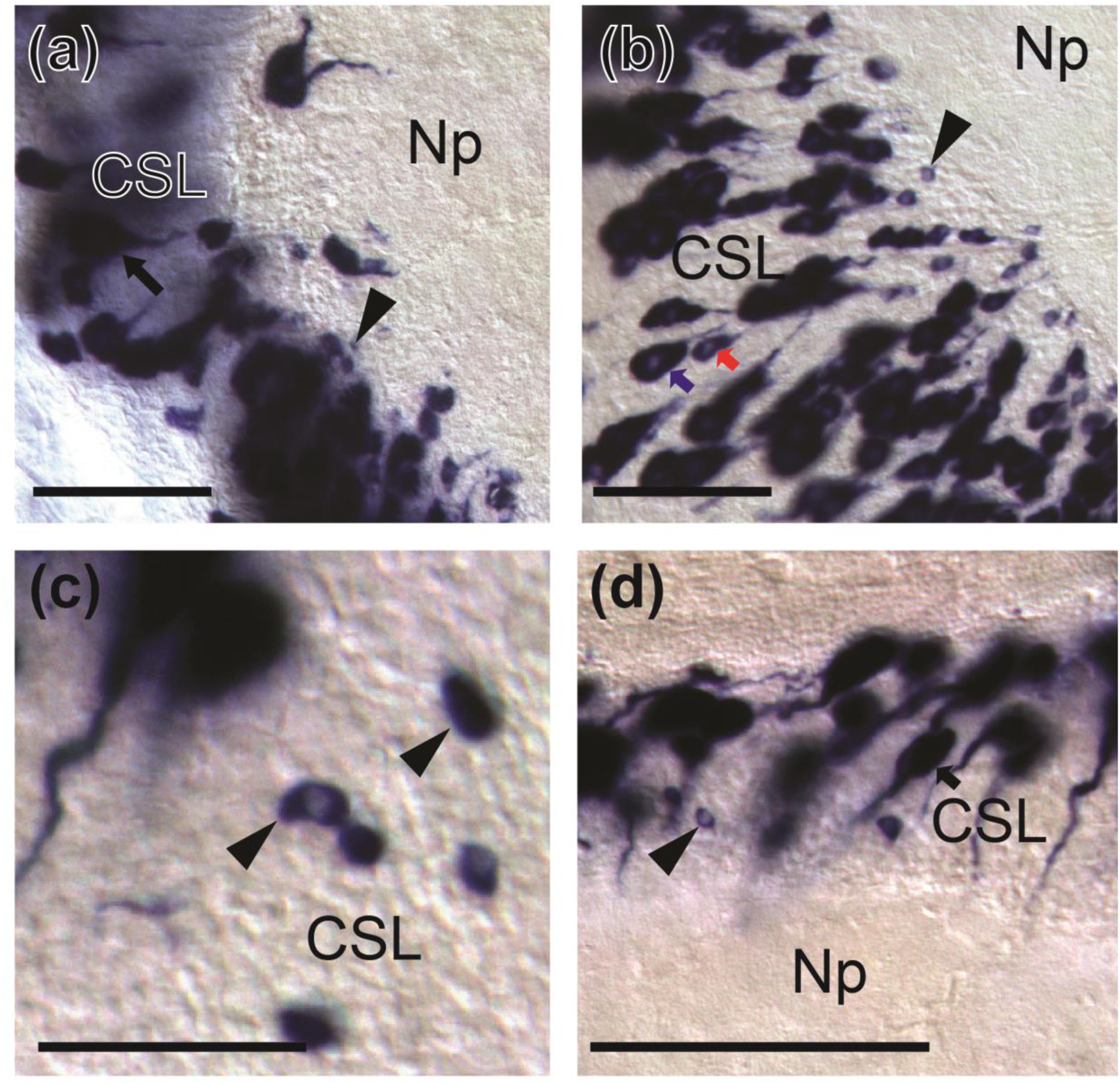
**Two morphologically distinct subtypes of neurons express neuropeptide Y precursor-encoding (pNPY) transcripts in the posterior buccal lobe (pBucL)**, as illustrated in (a) through (d). (a) and (b) are of coronal slices, and panels (c) and (d) are from sagittal slices. Arrows indicate pNPY neuron type eight, which possesses a large oval-shaped soma in the cell somata layer (CSL) with a projection directed toward the neuropil (Np). These neurons appear in a gradient of sizes ranging from about 10 to 35 µm in length (not including the projection). Examples of the variability in pNPY neuron type eight sizes are illustrated in (b), with a larger (~35 µm) cell soma indicated by a blue arrow, adjacent to a smaller (~20 µm) cell soma indicated by a red arrow. The second neuronal subtype identified in the pBucL are significantly smaller irregular ovals with no projections (pNPY neuronal type nine). These are primarily positioned at the edge of the cell somata layer proximal to the neuropil. A summary of the neuronal somata diversity (pNPY neuron types seven and eight) in this region can be found in Table 1. All scale bars measure 100 μm.

### pNPY expression in the Optic Lobe and Peduncle Complex is highly region-specific and sparse

Within the peduncle complex, located adjacent to the optic lobe on top of the optic tract, we observed a small cluster of pNPY positive neuronal somata in the olfactory lobe (Figure 5a, and 5b) (type 9 in Table 1). Specifically, this cluster of cells appears to be situated in the neuronal somata layer at the junction of the anterior and medial olfactory lobules (Figure 5a).

**Figure 5.**
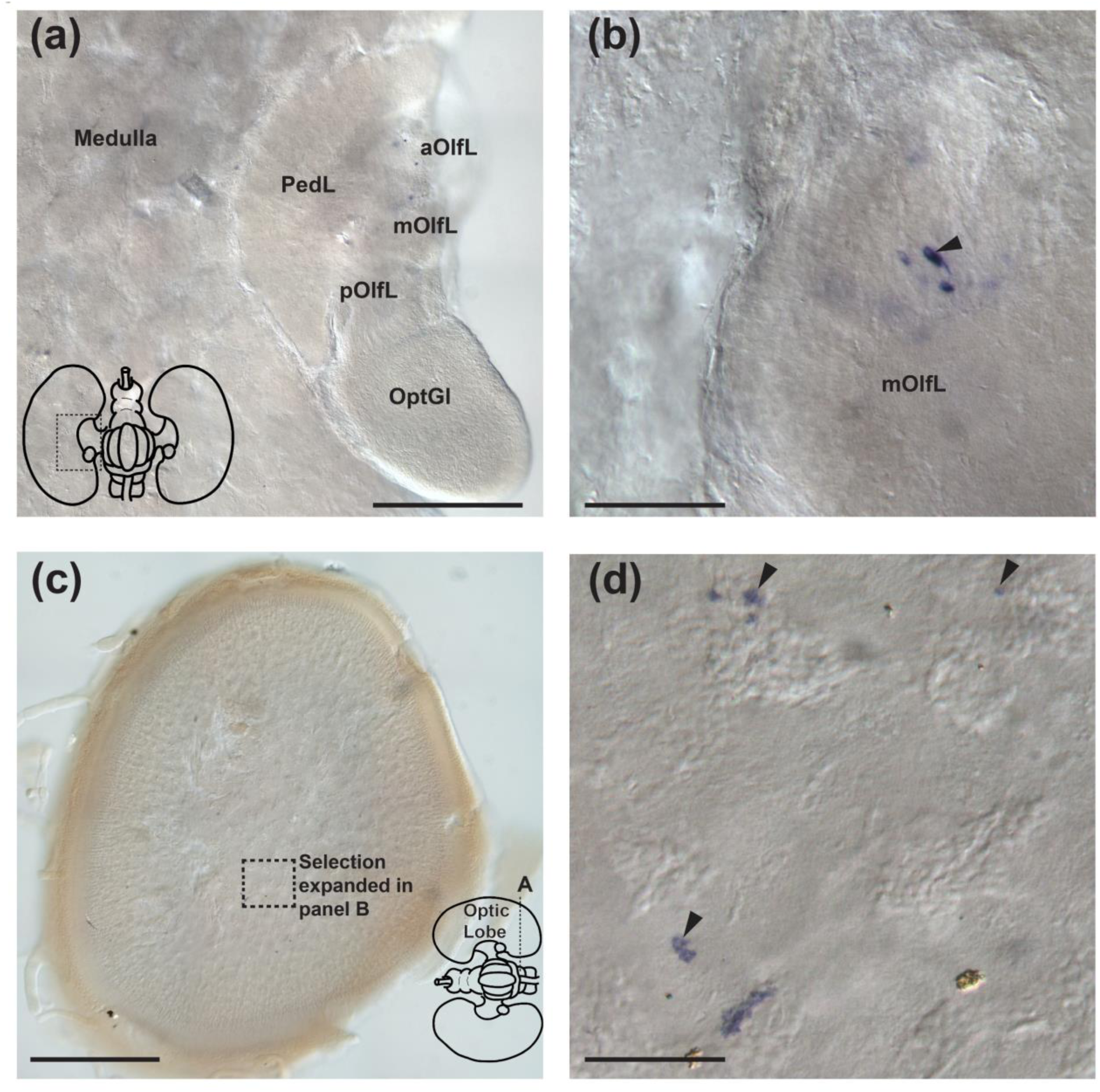
**A small cluster of pNPY expressing neurons is visible in the middle olfactory lobule,** but pNPY positive neurons of the Optic Lobe (OL) are scarce and not immediately apparent. Panel (a) illustrates the general location of pNPY positive neurons in the peduncle complex, specifically along the border of the anterior and middle lobule of the olfactory lobe (aOlfL and mOlfL respectively), which are anterior to the posterior olfactory lobule (pOlfL) and the optic gland (OptGL). (b) further magnifies this preparation to illustrate the elongated oval-shaped pNPY positive cells of the olfactory lobe, some of which have short visible processes. Panel (c) shows the overall infrequency of pNPY expressing neurons, as, at low magnification, the section appears relatively free of obvious labeling. Panel (d) is a further magnified view of the boxed section noted in (c). The arrowheads indicate tapered oval-shaped neuronal somata ranging from about 5 to 15 µm in diameter. A summary of the neuronal somata diversity (pNPY neuron type nine and ten) in this region can be found in Table 1. Scale bar measurements are (a and c) 500 μm, and (b and d) 50 μm.

Neurons expressing pNPY can be seen in some cell clusters of the optic lobe (Figure 5d) (type 10 in Table 1). These cells were scattered throughout the optic lobe. These cells share morphologically similarity with some of the NPY-IR optic lobe neuronal somata observed previously (Suzuki *et al.*, 2002). No pNPY positive neural processes are visible in the optic lobes.

## 4 Discussion

### NPY in molluscs

Detailed comparative analyses of the *Octopus* Neuropeptide Y precursor (Figure 1a) throughout metazoans builds on existing studies that have been primarily limited to the chordate lineage (Larhammar, 1996). Our analyses confirm that NPY may have originated in the common ancestor of all bilaterians. This conclusion is consistent with the computational cluster mapping (Jékely, 2013) of ancient neuropeptide families, indicating that the NPY protein family emerged in the Urbilateria. Our multiple sequence alignments reveal one mutation that is unique to the cephalopod lineage. Instead of the highly conserved dibasic cleavage site (KR) that follows the glycine (G) residue, an arginine (R) residue is found at position 78. In the cephalopods, this residue is followed by a serine (S) (or an aspartic acid (D) in *N. pompilius*). The conserved dibasic cleavage site is where posttranslational cleavage enzymes recognize and cut the prepropeptide before subsequent amidation. The variation here in cephalopod pNPY sequences may, therefore, indicate that the enzymes required to convert pNPY into bioactive NPY may be different from those employed in other closely related lineages. Although monobasic (as opposed to dibasic) cleavage sites are not uncommon in secretory molecules, their presence in cephalopod NPY sequences appears to be unique among molluscs.

An additional amino acid sequence variation can be seen at position 76 in Figure 1A. The N-terminal tyrosine (Y) at this site is the origin of the name “Neuropeptide Y” (Tatemoto K Fau - Carlquist *et al.*, 1982) in deuterostomes. However, in the case of many protostomes, like *Octopus*, the N-terminal amino acid is a phenylalanine (F). In some cases, like the insect *Drosophila melanogaster* (Fadda *et al.*, 2019), researchers have changed the name to the possibly more appropriate “Neuropeptide F”, but in this case, we will continue with the nomenclature of existing cephalopod NPY studies. Both phenylalanine and tyrosine possess hydrophobic side chains with the α-carboxyl group of phenylalanine being more acidic (1.83 vs. 2.2 pKa1) and its α-ammonium ion being very slightly less acidic (9.13 vs. 9.11 pKa2) than those of tyrosine. The only major structural difference between the two residues is that the phenyl group of tyrosine is hydroxylated, but that of phenylalanine is not. Despite minor structural and chemical differences in the N-terminal amino acids, functional studies across taxa show that this substitution at this site does not render the final product biologically inactive, nor does it dramatically change its physiological roles.

The alignments discussed above were used to generate a gene tree (Figure 1b) whose branches correspond with the predicted evolutionary history of particular animal lineages. This supports the scenario that pNPY sequences originally evolved in the common ancestor of protostomes and deuterostomes and subsequently radiated throughout characterized taxa without notable losses or duplications.

### pNPY expression in the *Octopus bimaculoides* brain reveals potential novel circuitry

The most striking expression pattern of pNPY was found in the dorsal basal lobe (DBL), where densely clustered neurons of the posterior DBL (pDBL) layer project anteriorly toward (and in some cases through) the anterior DBL (aDBL) as seen in Figure 2a. Although some pNPY positive neuronal somata (Figure 3c, cell type 4) are found in the aDBL, the majority of the DBL pNPY positive neurons originate in the pDBL (Figure 3a-b, cell types 1-3). According to the *Octopus* brain atlas (Young, 1971), the majority of the pDBL efferents project downward toward the median basal lobe (MBL), whose neuronal somata are also pNPY positive. In our study, however, the most obvious of the large pNPY positive tracts project anteriorly toward the anterior basal lobes (ABL), subfrontal lobes (SubFrL), and posterior buccal lobes (pBucL), or angle upward toward the subvertical lobe (SubVL).

NPY signaling between the pDBL and SubVL (part of the VL visual memory circuit) may occur in both directions, as we also saw pNPY positive neurons in the SubVL (Figure 3e) and bidirectional tracts between the SubVL and the pDBL have been described previously (Young, 1971). Based on their positions, a possible role for the pNPY positive projections between the pDBL and SubVL could be in processing or transmitting visual information, perhaps acting as an “integration” site between visual (VL circuit), tactile (SubFrL/ABL/pBucL), and ultimately motor centers.

To date, there are no described direct functional relationships between DBL neurons and the ABL, SubFrL, or pBucL, but our study has revealed clear evidence of pNPY projections originating in the DBL, terminating at or near each of these structures. The coronal brain cross-section in Figure 2d further verified that the long processes of pNPY positive DBL neurons were scattered throughout the basal lobes, indicating multiple targets from the same DBL region. The ABL has been implicated in the coordination of head and arms when handling prey (Young, 1971), and the pBucL and SubFrL are both key relay centers in the tactile learning and memory system, which is also imperative to prey capture. Currently, the only described inputs to the pBucL are the lateral inferior frontal (lIFL) and SubFrL (Young, 1971) of the tactile memory circuit and the axial nerve cords of the arms. Here we propose additional inputs from the long projections from the DBL, which may be using NPY as a signaling molecule to modulate the activity of other structures inextricably involved in memory and feeding behaviors (Di Cosmo *et al.*, 2014).

The abundant expression of pNPY encoding transcripts in both the large cells and the amacrine cells of the pBucL (Figure 2b-c and Figure 4) further suggests a role for NPY in this structure. Efferent tracts from the pBucL project back to the axial nerve cord and the SubFrL, so in this instance, NPY might be mediating a role for the pBucL as a second “integration center” between the tactile memory circuit and feeding behaviors elicited in the arms. Additional potential outputs for the pNPY positive neurons of the pDBL and MBL are the magnocellular and subesophageal lobes. Although there appears to be no pNPY expression in either of these locations, activities in these lobes may still be mediated by presynaptic pNPY positive neurons of the supraesophageal lobes.

Finally, the extrasomatic pNPY mRNAs that localized to some (but not all) neurites in the brain (Figures 2, 3, and 4) suggest that a select population of neurons transport mRNA along the axon, perhaps to synaptic regions (Puthanveettil *et al*., 2013). This may indicate a function of NPY transcripts in these areas, as these mRNAs could be locally translated and processed at the synapse instead requiring transport of synthesized protein down the length of the neurite after translation. The region-specific extrasomatic mRNAs may expedite fast and recoded responses specific to the highly regionalized brain structures, as was recently highlighted in squid (Vallecillo-Viejo *et al*., 2020).

As noted above, a role for NPY in food-related and motivational state learning and memory has also been characterized in larval neural circuits (Rohwedder *et al.*, 2015) and the mushroom bodies of adult (Krashes *et al.*, 2009) fruit flies, but one must use caution in making comparisons across such evolutionary distances. Despite the existence of sophisticated neuronal memory circuitry in flies and octopuses, these neural circuits very likely evolved independently of one another, thus requiring independent recruitment of any molecular commonalities (Moroz, 2009; Yoshida *et al*., 2015). Still, the possibility of parallel implementation of NPY for molecular mediation of memory circuitry across distant lineages suggests some unknown characteristic of NPY that facilitates its candidacy as a signaling molecule in memory circuit integration. Further examination of functional roles for NPY in intermediate species (those that punctuate the branches on the evolutionary tree) between flies and cephalopods will likely reveal whether these independent recruitment events resulted from some inherently advantageous and conserved aspect of NPY or are simply random.

### NPY expression in the *Octopus* brain and peduncle complex is functionally inconclusive

The entire DBL region of the *Octopus* brain receives input from the optic lobes via the optic tract, but projections from the olfactory lobe specifically target the pDBL and not the aDBL. It is possible that some of the large pNPY positive tracts that originate in the pDBL project back toward the ipsilateral middle olfactory lobule (Messenger, 1967). Experimental studies have indeed shown that degeneration of the pDBL neurons induces optic gland enlargement and the onset of reproductive processes (Wells *et al.*, 1959), suggesting an integral role for pDBL neurons in the regulation of reproduction. Given the generally conserved role for NPY in feeding and the apparent interplay between the pDBL and the optic gland via the olfactory lobe, it is possible that the abundant pNPY transcript expression in the pDBL and efferent tracts might be involved in maintaining feeding behaviors while blocking the animal’s transition to reproductive life stages in the olfactory lobe as suggested previously (Di Cristo *et al.*, 2017).

Based on existing models for the role of NPY in *Octopus* (Di Cristo *et al.*, 2017), we predicted that pNPY transcripts would be expressed in the olfactory lobe of the juvenile adult male octopuses used for this study. We consistently identified a small cluster of pNPY-positive neurons at the junction of the middle and anterior olfactory lobules (Figure 5 a and b). The neuronal somata expressing pNPY in the olfactory lobe appear to be elongated ovals with projections pointed toward the neuropil of the middle olfactory lobule. These pNPY positive cell clusters were quite small and lacked the dramatically abundant expression seen in the central brain tissues. However, it is imprudent to assume insignificance of the small cluster of pNPY expressing cells just because they are visually underwhelming. For example, there are only 20 or 26 (in female vs. male respectively) NPY/NPF expressing neurons in the entire nervous system of *D. melanogaster*, but a myriad of functions have been attributed to this small pool cells (Nässel *et al.*, 2011). We are not suggesting that the small pool of NPY/NPF expressing neurons in flies bears any homology to any of the pNPY expressing cell populations of the *Octopus* brain, but highlighting the fact that in some cases, the number of cells involved in a physiological process does not directly correlate with the scope of their function. Our detection of a small cluster of pNPY mRNA expressing neurons in the olfactory lobes are therefore not inconsistent with the existing hypothesis that NPY may act as a signaling molecule among the olfactory lobes, optic glands, and the brain (Di Cosmo and Polese 2014) (via the olfactory nerve (Young, 1971)).

Interestingly, the expression patterns of pNPY mRNAs were different from that of NPY protein localization studies using immunohistochemistry (Suzuki *et al.*, 2002). NPY-immunoreactive (NPY-IR) cell somata and fibers were previously identified throughout the whole peduncle and optic lobe complexes, including in the optic glands. Localization patterns of transcripts encoding pNPY detected by *in situ* hybridization were restricted to the middle olfactory lobe cluster mentioned above. Another noted inconsistency between our *in situ* hybridization mapping and NPY immunohistochemical studies (Suzuki *et al.*, 2002) is the stark difference between the distribution of NPY positive cells and fibers in the optic lobes (Figure 5c and d). This could be due to the fact that in this study, we used only juvenile *O. bimaculoides* males. However, one might expect to see different NPY expression in the optic lobes of immature females, similarly to the NPY-IR expression previously reported (Suzuki *et al*., 2002).

There are further possible explanations for the NPY distribution discrepancy across studies. Because *in situ* hybridization only reveals mRNA localization, far more structures (mainly neuronal processes) were detected using immunohistochemistry (Suzuki *et al.*, 2002) than by our *in situ* hybridization methods. Therefore, the discrepancy could be somewhat attributed to the limitations of each technique. pNPY or NPY protein might be very stable, negating a need for constant mRNA production and storage throughout the necessary tissues. This explanation, however, does not fit with the abundant expression of pNPY mRNA in the brain, which indicates that pNPY mRNA exists at relatively high levels in specific lobes. An additional possibility for the discrepancies in the distribution of pNPY in this study and NPY-IR cells in previous studies (Suzuki *et al.*, 2002) is the noted cross-reactivity of their antibody with the related peptide, Peptide YY (P-YY). In this case, our study provides a method to positively distinguish the P-YY neurons from the true pNPY expressing neurons of the *Octopus* nervous system.

## 5 Conclusions

Here we characterized the expression of the Neuropeptide Y prohormone in *Octopus bimaculoides* nervous system (an important reference species in evolutionary neuroscience (Albertin *et al*., 2012)), adding novel molecular and structural insights into the mechanisms controlling feeding and reproduction. We noted a generally conserved putative active site, flanked by a minor lineage-specific sequence variation at the N-terminal cleavage site in *Octopus* pNPY.

We have also identified the distinct expression of pNPY in specialized regions of the brain, including two potential “integration centers” where visual, tactile, and behavioral neural circuitry converge: the dorsal basal lobe and the posterior buccal lobe. Each of these possible integration centers can produce pNPY as the signaling molecule for communications between the olfactory lobe (which controls reproductive behaviors), subvertical and subfrontal lobes (employed in visual and tactile learning and memory respectively), and the anterior basal lobes and axial nerve cords (both implicated in feeding behaviors). These centers and their accessory structures may control feeding and reproductive behaviors, using NPY as the signaling peptide with multiple integrative functions. The extrasomatic localization of pNPY mRNA may add an additional layer of temporal regulation of the complex behaviors in cephalopods.

The partial overlap of immunohistochemical and *in situ* hybridization localization provides intriguing bases for functional and temporal hypotheses about the role of NPY in *Octopus* nervous systems. If the current models (Di Cosmo and Polese 2014) are accurate, then it would also follow that the pNPY expressing neurons of the olfactory lobe exert an inhibitory effect on the optic gland and the onset of reproductive behaviors. NPY synthesis and *in-vivo* administration (or manipulation of the predicted NPY receptor) and subsequent behavioral and physiological characterization would help us to understand the exact regulatory role of NPY in *Octopus*. Behavioral and physiological characterization of *Octopus* after exogenous application of NPY would help us determine whether the primary function of NPY expressing neurons is to regulate a component of the memory circuitry, mediate the transition between feeding and reproduction life stages, or/and something yet un-proposed, suggesting a possible pleiotropic role played by NPY in different regions of the nervous system.

## Data Availability

The data that support the findings of this study are available from the corresponding author upon reasonable request.

## Author contributions

G.C.W. and L.L.M.: Conceptualization, Data curation, Formal analysis, Investigation, Methodology, Software, Validation, Visualization, Writing original draft, Writing-review & editing. L.L.M.: Funding Acquisition, Project Administration, Resources, and Supervision. G.P. and A.DC. Validation, Writing-review & editing

## Acknowledgments

The authors would like to thank Yelena Bobkova and Tatiana Moroz for technical laboratory assistance, and Dr. Caleb Bostwick for editing. This work was supported by the Human Frontiers Science Program (RGP0060/2017) and National Science Foundation (grants 1146575, 1557923, 1548121 and 1645219 to L.L.M); we also thank the Compagnia di San Paolo for supporting this study by a “Single Center Research Grant in Neuroscience” (Protocol 29-11 A.DC).

## Footnotes

1 It is important to note that the term “olfactory lobe” was designated to this structure because of its direct anatomical connection to an olfactory organ, and does not imply that the only function of the lobe is “olfactory” or that all neurons housed here play a direct role in chemosensation. The specific function(s) of this lobe are likely diverse and still being elucidated.

